# Evolution of the quorum sensing systems in *Pseudomonas aeruginosa* can involve both loss of regulon function and network modulation

**DOI:** 10.1101/2022.04.27.489656

**Authors:** Priyanikha Jayakumar, Alexandre R. T. Figueiredo, Rolf Kümmerli

## Abstract

*Pseudomonas aeruginosa* populations evolving in cystic fibrosis (CF) lungs, animal infection models, natural environments or *in vitro* undergo extensive genetic adaption and diversification. A common mutational target is the quorum sensing (QS) regulon, a three-unit regulatory system that controls the expression of a suite of virulence factors and secreted public goods. Three scenarios have been advocated to explain selection for QS mutants, which include (I) disuse of the regulon, (II) cheating on public goods, or (III) modulation of the network. Here, we test these scenarios by examining a set of 61 QS mutants from an experimental evolution study. We observed non-synonymous mutations in all three QS systems – Las, Rhl and PQS. Most Las mutants carried large deletions, resulting in loss of QS function, and the inability to produce QS-regulated traits (scenario I or II). Conversely, phenotypic and gene expression analyses of Rhl mutants support network modulation (scenario III), as these mutants overexpressed the Las and Rhl regulators and showed an altered QS-regulated trait production portfolio. PQS mutants also showed patterns of network modulation (scenario III), spurring strain diversification and phenotypic trade-offs, where the upregulation of certain QS traits is associated with the downregulation of others. Overall, our results indicate that mutations in different QS systems lead to diverging effects on the social portfolio of bacterial populations. These mutations might not only affect the plasticity and diversity of evolved populations but could also impact bacterial fitness and virulence in infections.

**Importance:** *Pseudomonas aeruginosa* uses quorum sensing (QS), a three-unit multi-layered network, to coordinate expression of traits for growth and virulence in the context of infections. Despite its importance for bacterial fitness, the QS regulon appears to be a common mutational target during long-term adaptation of *P. aeruginosa* in the host, natural environments and experimental evolutions. This raises the questions why such an important regulatory system is under selection and how mutations change the portfolio of QS-regulated traits. Here, we examine a set of 61 naturally evolved mutants to address these questions. We found that mutations involving the master regulator, LasR, resulted in an almost complete breakdown of QS, while mutations in RhlR and PqsR resulted in modulations of the QS regulon, where both the QS regulon structure and the QS-regulated trait portfolio changed. Our work reveals that natural selection drives diversification in QS activity patterns in evolving populations.

## Introduction

*Pseudomonas aeruginosa* is an opportunistic bacterial pathogen responsible for chronic infections, especially in individuals with the genetic disorder cystic fibrosis (CF) (Koch and Hoiby, 1993; Parkins, Somayaji and Waters, 2018). *P. aeruginosa* lineages isolated from patients are often characterized by a series of specific mutations, which have been traditionally interpreted as adaptations to the CF lung environment (Smith *et al*., 2006; Folkesson *et al*., 2012; Dettman *et al*., 2013). The quorum-sensing (QS) regulon is one of the commonly observed mutational hotspots (Schaber *et al*., 2004; Smith *et al*., 2006; Damkiær *et al*., 2013; Marvig *et al*., 2015; Feltner *et al*., 2016; Winstanley, O’Brien and Brockhurst, 2016). As *P. aeruginosa* uses QS to regulate a suite of virulence factors (Rumbaugh *et al*., 1999; Pearson *et al*., 2000; Lesprit *et al*., 2003), it is rather surprising to see that mutations in QS regulators (often interpreted as loss-of-function mutations) are favored in an infectious context, where virulence factors are important.

QS signaling in *P. aeruginosa* is mediated by two N-acyl homoserine lactone (AHL)-dependent QS systems, the Las and Rhl systems, as well as the *Pseudomonas* Quinolone Signal (PQS) system (Williams and Cámara, 2009; Nadal Jimenez *et al*., 2012; Lee and Zhang, 2015). Each system synthesizes its own signal (Las: 3O-C12-HSL; Rhl: C4-HSL, PQS: 2-heptyl-3-hydroxy-4-quinolone) that binds to its cognate receptor (LasR, RhlR and PqsR). Signal-receptor complexes form transcriptional regulators that control the expression of a suite of virulence factors, including a set of secreted proteases, biosurfactants, toxins, and biofilm formation. The induction of these virulence factors depends on surpassing a signal threshold concentration, which is often reached at high bacterial population densities. The QS systems are arranged in a hierarchical signaling cascade where the Las system positively regulates both the Rhl and PQS systems through the Las signal-receptor dimer complex. The PQS system also positively regulates the Rhl system, but Rhl in turn inhibits the PQS system (Diggle *et al*., 2007; Köhler, Buckling and Van Delden, 2009).

*P. aeruginosa* isolates from chronically infected CF patient lungs frequently contain mutations in the master transcriptional regulator, LasR, thereby influencing the activity of all three QS systems (Feltner *et al*., 2016; Chen *et al*., 2019). While initially interpreted as a specific adaptation to the CF lung environment, it has become clear that *lasR* mutants are also selected for under many other conditions, including chronic wounds (Vanderwoude *et al*., 2020), corneal infections (Preston *et al*., 1997; Hammond *et al*., 2016), ventilator-associated pneumonia (Köhler, Buckling and Van Delden, 2009), infections in *Caenorhabditis elegans* (Jansen *et al*., 2015; Granato *et al*., 2018), as well as in the absence of a host such as in natural environments (Groleau *et al*., 2021) and in experimental evolutions (Wilder, Diggle and Schuster, 2011; Kostylev *et al*., 2019; Scribner *et al*., 2021; Smalley *et al*., 2022). Why then, are *lasR* mutants consistently favored across these difference environments? Three competing hypotheses have been advocated. First, mutations in *lasR* lead to a loss of function in QS-regulated phenotypes and are favored because QS is no longer needed in the respective environments, especially during growth in rich medium (D’Argenio *et al*., 2007). Second, QS is still required but *lasR* mutants are cheaters that no longer respond to the QS signal. They refrain from producing QS-regulated traits, yet, still benefit from the shared pool of QS-regulated traits in the environment (proteases, biosurfactants, toxins) produced by QS wild type cells (Diggle *et al*., 2007; Köhler, Buckling and Van Delden, 2009; Rumbaugh *et al*., 2009). Third, mutations in *lasR* may modulate the QS regulon itself by either changing its sensitivity or remodeling the hierarchal network as an adaptation to the prevailing conditions (Chen *et al*., 2019; Kostylev *et al*., 2019). This hypothesis has been fueled by recent findings that evolved *lasR* mutants have diverse phenotypes and are not necessarily null mutants (Jansen *et al*., 2015; Feltner *et al*., 2016; Cruz *et al*., 2020).

To obtain a deeper understanding of how mutations in the QS regulon affect downstream phenotypes and QS network topology, we used a set of 61 experimentally evolved QS mutants to investigate whether these mutants have lost the ability to produce QS-regulated traits (supporting either the first or the second hypotheses) or if they show an altered QS-regulated trait expression portfolio (supporting the third hypothesis). The mutant collection stems from an experimental evolution study performed in our laboratory that focused on the evolution of iron uptake systems under various *in vitro* conditions (Figueiredo, Wagner and Kümmerli, 2021). The experiment was initiated with *P. aeruginosa* PAO1 wild type populations and ran for 200 consecutive days. Although the QS regulon was not the focus of this work, whole-genome sequencing of evolved clones revealed an accumulation of non-synonymous mutations in all three QS systems, corroborating the notion that mutations in QS systems are commonly favored in this species. In a first step, we conducted an in-depth genomic analysis on the types, size and location of mutations found in the Las, Rhl and PQS systems. Next, we screened all mutants for four QS-regulated traits to examine which type of mutations lead to a loss of function versus a modulated QS response. The four QS-regulated traits are (i) proteases, used to digest extracellular proteins, (ii) pyocyanin, a broad-spectrum toxin, (iii) rhamnolipid biosurfactants, for group-level motility and (iv) the ability to form surface-attached biofilms. Finally, we picked a subset of QS mutants with apparent QS-regulon modifications and investigated whether these mutations alter the gene expression of QS-regulators and downstream regulated traits.

## Materials and methods

### Bacterial strains

We analyzed a collection of 61 experimentally evolved *P. aeruginosa* clones from Figueiredo, Wagner and Kümmerli (2021) (Table S1). All clones have a common ancestor, the PAO1 wild type strain (ATCC 15692). We grouped the evolved clones based on the mutations accumulated in either a single (Las, Rhl or PQS systems), or in multiple QS systems. For the growth and QS-phenotype screening assays, we further used the ancestral PAO1 wild type strain and three isogenic QS mutants constructed from the same PAO1 background. The isogenic QS mutants are deficient in the production of either one of two QS receptors, LasR (*ΔlasR*), RhlR (*ΔrhlR*), or both receptors (*ΔlasR*-*ΔrhlR*). These are loss of function mutants and were used as controls for the screening of QS-regulated trait production.

To be able to track gene expression in a subset of mutated clones (n = 5), we engineered double fluorescent transcriptional reporter fusions to measure the simultaneous expression of (1) *lasR-gfp* and *rhlR-mCherry,* and (2) *lasB-gfp* and *rhlA-mCherry*. A single copy of the double reporter construct was chromosomally integrated in the experimentally evolved clones at the neutral attTn7 site using the mini-Tn7 system (Choi and Schweizer, 2006). Detailed step-by-step cloning protocol is described elsewhere (Jayakumar et al., 2021). We used *Escherichia coli* CC118 λpir for all intermediary steps in our cloning work (see Table S2 for a full list of non-experimentally evolved strains and plasmids used).

### Experimental evolution

The protocol of the experimental evolution study is described in detail elsewhere (Figueiredo, Wagner and Kümmerli, 2021). Briefly, experimental cultures were initiated with ancestral PAO1, and evolving populations were propagated for 200 consecutive days, during which approximately 1,200 generations occurred. Evolving populations were cultured in casamino acid (CAA) medium (5 g/L casamino acids, 1.18 g/L K_2_HPO_4_.3H_2_O, 0.25 g/L MgSO_4_.7H_2_O, 25 mm HEPES buffer) with varying iron availabilities (no FeCl_3_ added, 2 µm FeCl_3_, or 20 µm FeCl_3_ to achieve conditions of low, intermediate and high iron availability, respectively) and environmental viscosities (0%, 0.1% or 0.2% [weight/volume] agar to represent low, mid or high spatial structure, respectively). While these environmental conditions were important for the initial study design, they do not serve purpose for the current study, as QS mutants arose in all nine environments.

### Bioinformatical analysis

Figueiredo et al. (2021) sequenced the whole genome of 119 evolved clones, among which 61 (51.2%) had mutations in genes of the QS regulons. Experimentally evolved clones were sequenced on the Illumina NovaSeq6000 platform (paired-end, 150 base-pair reads). Single nucleotide polymorphisms (SNPs) and microindels (small insertions and deletions) were detected by aligning the obtained reads to the *P. aeruginosa* PAO1 reference genome with the BWA “mem” algorithm followed by variant-calling with BCFTOOLS and annotation with SNPEFF. Large deletions and duplications were detected with CLC Genomics Workbench. Details on the bioinformatic analysis are described in Figueiredo, Wagner and Kümmerli (2021).

To map the position of SNPs and microindels within each QS gene, we compared the sequenced genome of the single evolved clones to the *P. aeruginosa* PAO1 reference genome on www.pseudomonas.com. Using published protein database of the QS signal-receptor complexes on InterProScan, we further obtained the classification of protein families and domains and extracted the information on the amino acid residues of the ligand- and DNA-binding domains of the Las, Rhl and PQS transcriptional regulator complexes. Finally, to evaluate mutational hotspots, we mapped the position of the evolved mutations to the reference gene sequence of *lasR, rhlR* and *pqsR*.

### Growth measurements

For all experiments, we pre-cultured single clones from freezer stocks in 6 ml Lysogeny Broth (LB), at 37°C, 220 rpm for 18 hours. Prior to experiments, we washed overnight cultures twice with 0.8% NaCl and adjusted to an optical density at 600 nm (OD_600_) of 1. To measure growth, we inoculated cells from overnight pre-cultures into fresh 1.5 mL LB medium to a final starting OD_600_ of 0.01 in 24-well plates and incubated them at 37°C for 24 hours under shaken conditions (170rpm). The reasoning of this experiment was to obtain a proxy for fitness for all evolved clones relative to the ancestor in a standard medium, where the QS network is induced, but not essential (Jayakumar *et al*., 2021). After 24 hours, we measured growth as OD_600_ in a microplate reader (Tecan Infinite M-200, Switzerland).

### Pyocyanin production

To measure the production of pyocyanin, we collected the bacterial cultures after 24h of growth in LB medium (described above) in 2 mL reaction tubes. We thoroughly vortexed, and centrifuged them at 12,000 g for 10 minutes to pellet bacterial cells. We then transferred the cell-free supernatants to fresh 2 mL reaction tubes. For each clone, we transferred four aliquots of 200 µL of the cell-free supernatant to 96-well plates, and quantified pyocyanin by measuring optical density at 691 nm in a microplate reader. LB medium was used as a blank control.

### Rhamnolipid production via drop collapse assay

We used the drop collapse assay to measure the production of rhamnolipids. We collected cell-free supernatant of bacterial cultures grown in LB medium as described above. For each clone, we plated 5 µL of the cell-free supernatant on the lids of 96-well plates and measured the droplet surface area after one minute (Kramer, López Carrasco and Kümmerli, 2020). Surface tension decreases with increasing concentrations of biosurfactant in the supernatant, therefore resulting in the collapse of droplets (Bodour and Miller-Maier, 1998). We took pictures of the lids and measured droplet surface area with the Image Analysis Software *ImageJ.* LB medium was used as a blank control. To quantify biosurfactant production based on droplet surface area, we made a calibration curve with a known range of synthetic rhamnolipid (Sigma-Aldrich, Switzerland) concentrations (ranging from 0-0.2 g/L) and measured their respective droplet surface area.

### Protease production

We used the azocasein assay to measure protease production. For this, we inoculated cells from overnight pre-cultures into 1.5 mL casein medium (5 g/L casein from bovine milk, 1.18 g/L K_2_HPO_4_.3H_2_O, 0.25 g/L MgSO_4_.7H_2_O) to a final starting OD_600_ of 0.01 in 24-well plates, and incubated cultures at 37°C for 48 hours under shaken conditions (170rpm). After 48 hours, we transferred the bacterial cultures to 2 mL reaction tubes, vortexed thoroughly, and centrifuged at 12,000 g for 10 minutes to pellet bacterial cells. Next, we transferred the cell-free supernatants to fresh 2 mL reaction tubes. We first treated aliquots of 40 µL cell-free supernatants with 120 µl phosphate buffer (50 Mm, pH ≈ 7.5) and 40 µL azocasein (30 mg/mL), and subsequently incubated them at 37°C for 30 minutes. We stopped the reaction with 200 µL trichloroacetic acid (20 %). We centrifuged treated supernatants at 12,000 g for 10 minutes and collected and transferred fresh supernatants into new 96-well plates. We quantified protease production as optical density at 366 nm in a microplate reader. Casein medium treated with azocasein was used as a blank control. All media components were purchased from Sigma-Aldrich, Switzerland.

### Biofilm measurements

We used the crystal violet assay to measure the ability of evolved clones to form surface-attached biofilms. We prepared overnight pre-cultures of single clones from freezer stocks in 200 µL LB medium in 96-well plates and incubated them at 37°C under static condition for 24 hours. We measured the growth of pre-cultures at OD_600_ using a microplate reader. Then, we diluted the pre-cultures to a starting OD_600_ of 0.01 in fresh 100 µL LB medium in a 96-well round bottom plate (No. 83.3925.500, Sarstedt, Germany) and incubated at 37°C under static conditions for 24 hours. Subsequently, we carefully transferred the cultures to a fresh flat-bottom 96-well plate and measured growth at OD_600_ in a microplate reader. We added 100 µL of 0.1% crystal violet to each well of the round bottom plate to stain the surface-attached biofilm and incubated the plates at room temperature for 30 minutes. Then, we carefully washed the wells twice with ddH_2_0 to remove the crystal violet solution and left them to dry at room temperature for 15 minutes. Next, we added 120 µL of dimethyl sulfoxide (DMSO) to each well to solubilize the stained biofilm and incubated the reaction at room temperature for 20 minutes. Finally, we measured optical density at 570 nm in a microplate reader, and the production of surface-attached biofilm was quantified by calculating the “Biofilm Index” (OD_570_ / OD_600_) for each well (Savoia and Zucca, 2007). LB medium treated with crystal violet and DMSO was used as a blank control.

### Gene expression measurement

We inoculated fluorescent gene reporter cells from overnight cultures into fresh LB medium to a final starting OD_600_ of 0.01 in individual wells on 96-well plates. Plates were incubated at 37°C in a microplate reader. We measured mCherry fluorescence (excitation: 582 nm, emission: 620 nm), GFP fluorescence (excitation: 488 nm, emission: 520 nm) and growth (OD_600_) every 15 minutes (after a shaking event of 15 seconds) over a duration of 24 hours. To remove background fluorescence, we measured the mean fluorescence intensity of the untagged PAO1 wild type strain in the mCherry and the GFP channels across time and subtracted these values from the measured mCherry and GFP fluorescence values of the QS gene reporter strains at each time point.

### Statistical analysis

We performed all statistical analyses with R studio (version 3.6.1). For all datasets, we consulted Q-Q plots and the Shapiro-Wilk test to examine whether the residuals were normally distributed. We used one-way ANOVA and post-hoc Tukey’s HSD to compare growth and QS-regulated traits between the different mutant categories, and between the mutant categories and the ancestral wild type. We performed a principal component analysis (PCA) on the clonal phenotypes using the vegan package in R (version 2.5-7) (Oksanen *et al*., 2020). We further tested whether mutant categories differ in their evolved QS trait profile using permutational multivariate analysis of variance (PERMANOVA). To compare gene expression trajectories, we fitted a parametric growth model (logistic model) in R and extracted the area under the curve (AUC) of each clone. Then, we used one-way ANOVA to compare the AUC between the mutant categories.

## Results

### Mutational patterns across the three QS regulons of *P. aeruginosa*

Among the 61 evolved clones, we found 68 mutations in genes of the three QS regulon (see Table 1 for an overview and Table S1 for individual clones). We detected 29 large-scale deletions (>4,903 bp), 30 single nucleotide polymorphisms (SNPs), and 9 microindels representing small deletions (max 12 bp) in the genes within the Las, Rhl and PQS systems. Most mutations were observed within the Las system (n=35), followed by the PQS (n=28) and Rhl (n=5) systems.

**Table 1:** Mutations within each QS system.

| System | Gene | Description | Number of clones |
| --- | --- | --- | --- |
| Las | *lasI-rsaL-lasR* | Las signal, repressor and receptor | 29 |
|  | *lasR* | Las receptor | 6 |
| Rhl | *rhlR* | Rhl receptor | 5 |
| PQS | *pqsABCD* | PQS signal operon | 3 |
|  | *pqsE* | PQS signal operon | 4 |
|  | *pqsR* | PQS receptor | 21 |

#### Mutations within the Las regulon

The majority of mutations in the Las regulon entail large-scale deletions (n = 29, 82.9%, ranging from 4,903 bp to 65,969 bp), where the Las signal synthase (*lasI*), the negative repressor (*rsaL*) and the Las receptor (*lasR*) were deleted, in addition to other genes. The exact position and size of the deletions are shown in Fig. 1A. In contrast, we only found a small number of SNPs (n = 6, 17.1%) in the *lasR* receptor, of which five are located in the same region of the DNA binding domain (Fig. 1B). The single mutant that has a SNP at a different location also has a mutation in the PQS system.

**Figure 1:**
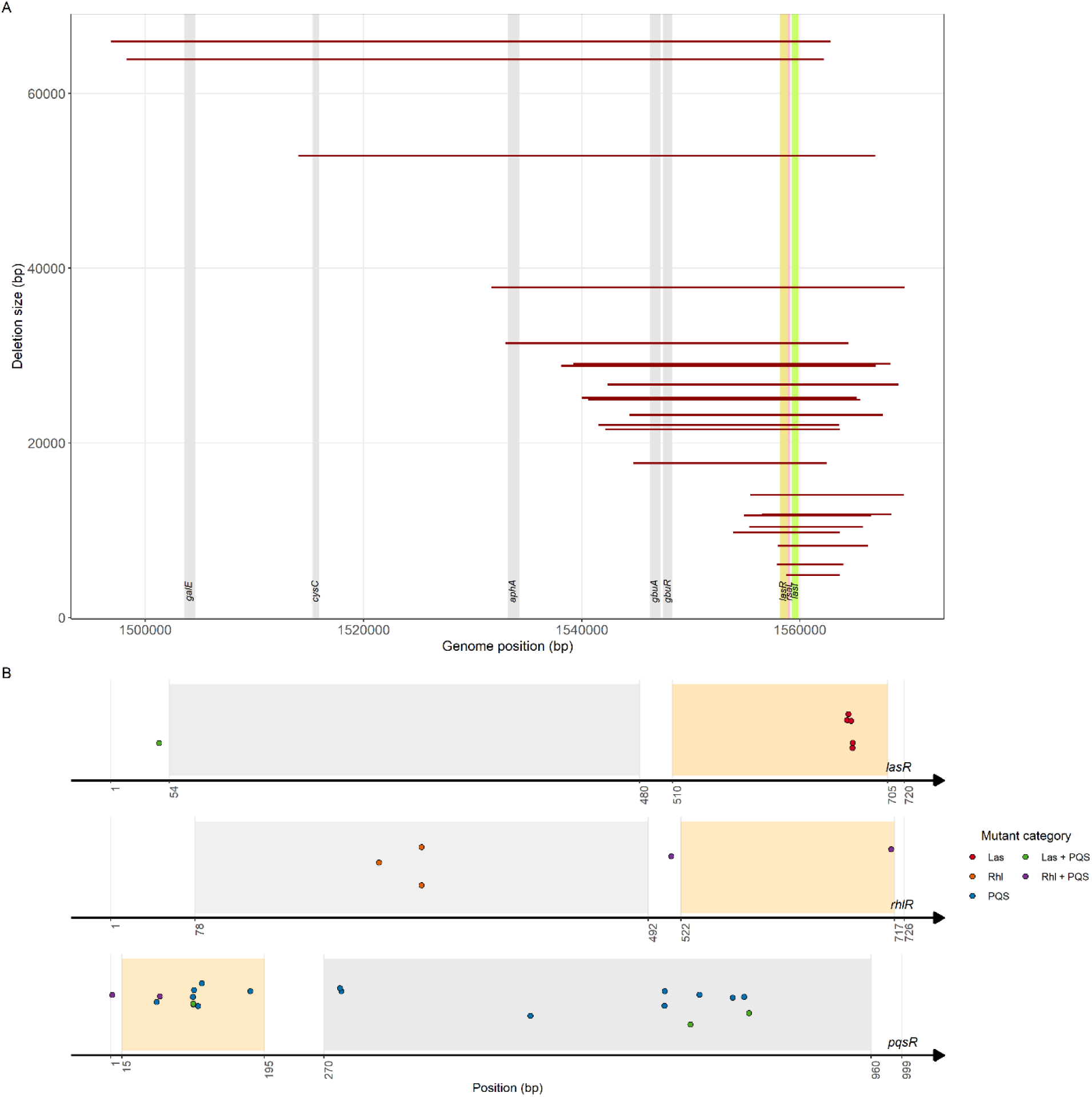
Experimentally evolved mutations in the QS systems of *P. aeruginosa*. (A) Size and position of large-scale deletions that include *lasI, rsaL* and *lasR* of the Las system. Shaded area represents position of genes in the genome. (B) Evolved mutations (SNPs and microindels) in the genes encoding the receptors of Las (*lasR -* 720 bp), Rhl (*rhlR -* 726bp) and PQS (*pqsR -* 999 bp) systems. Each dot represents the position of the mutated nucleotide within each gene. Grey and orange areas represent the ligand (i.e., signal)-binding and the DNA-binding domains, respectively.

#### Mutations within the Rhl regulon

In total, we found five SNPs in the gene coding for the Rhl receptor (*rhlR*). Two of the five mutants also have SNPs in the PQS system. Although the numbers are too few to obtain a conclusive pattern on the location of mutations, we found that the three clones that only had *rhlR* mutated all have SNPs in the ligand-binding site of *rhlR* (Fig. 1B). Meanwhile, the two *rhlR*-PQS double mutants have their SNPs outside the ligand-binding site.

#### Mutations within the PQS regulon

Out of the 26 clones with mutations in the PQS system, 19 have single mutations within the PQS regulon, 2 clones have double mutations within the PQS regulon, while 5 clones share one other mutation in either the Las or the Rhl system. Altogether, there were 28 mutations, comprising of 2 SNPs and 1 microindel in the PQS signal operon (*pqsABCD*), 4 SNPs in the *pqsE* gene and 13 SNPs and 8 microindels in the gene encoding the PQS receptor (*pqsR*). When mapping the mutations in *pqsR*, we found that the SNPs and microindels occurred both in the DNA- and the ligand-binding domains (Fig. 1B).

### QS system-specific mutations drive divergence in the production of QS-regulated traits

Next, we explored how mutations in the Las, Rhl and PQS regulons link to growth and QS trait expression (proteases, rhamnolipids, pyocyanin and biofilm). We grouped mutants into five categories: (i) clones with mutations in the Las receptor, *lasR,* and large-scale Las deletions (these two classes were combined because there was no difference in their phenotypes); (ii) clones with mutations in the Rhl regulon alone; (iii) clones with mutations in the PQS regulon alone; (iv) clones with mutations in the Las and the PQS regulons; (v) clones with mutations in the Rhl and the PQS regulons. For the statistical analysis, we further included the ancestral wild type as sixth category and compared whether there are significant differences in growth and QS-regulated trait production between the mutant categories and the wild type, as well as between the five mutant categories.

Our growth assay in LB medium revealed no significant difference in endpoint growth between any of the five mutant categories and the ancestral wild type (Fig. 2A, one-way ANOVA, F_5,177_ = 0.587, p = 0.710). However, there were considerable differences in growth performance between evolved clones within certain mutant categories, especially among those with mutations in the Las regulon.

**Figure 2:**
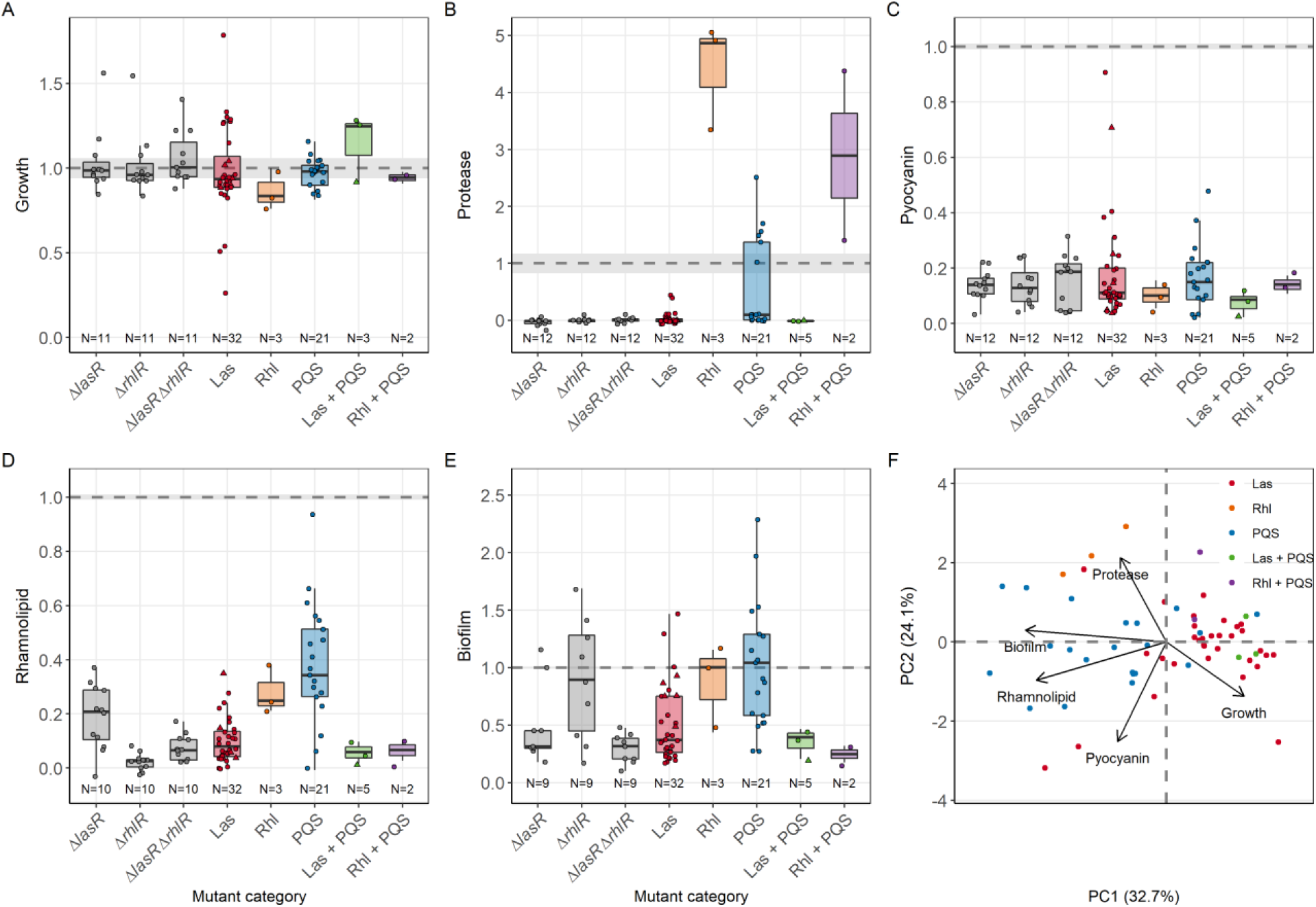
Phenotypic profile of experimentally evolved *P. aeruginosa* isolates with mutations in the QS regulon. (A) Endpoint planktonic growth (OD at 600 nm) in LB medium after 24 hours, and production of four QS-regulated traits: (B) protease, (C) pyocyanin, (D) rhamnolipid and (E) surface-attached biofilm across clones with mutations in either a single (Las, Rhl or PQS systems) or in multiple QS systems (Las + PQS, and Rhl + PQS). All values are expressed relative to the corresponding value of the ancestral PAO1 wild type strain (mean ± standard error indicated as dotted lines and shaded areas, respectively). Lab-generated QS mutants deficient in the production of either one of the two QS receptors, LasR (*ΔlasR*), RhlR (*ΔrhlR*), or both receptors (*ΔlasR*-*ΔrhlR*) were used as controls for the production of QS-regulated traits in loss-of-function mutants. (F) Principal component analysis (PCA) on growth and the production of the four QS-regulated traits. Each data point represents the average of at least three independent replicates.

For proteases, we found significant differences in the production levels across the five mutant categories and the ancestral wild type (Fig. 2B, one-way ANOVA, F_5,156_ = 20.388, p < 0.001). All clones with mutations in the Las system (including the Las-PQS double mutants) had lower protease production compared to the ancestral wild type, with most clones having almost completely abolished production, similar to the lab-generated *lasR* mutant. Meanwhile, all clones with mutations in the Rhl system (including the Rhl-PQS double mutants) produce higher amounts of proteases than the ancestral wild type. This observation is diametrically opposite to the pattern seen in the lab-generated *rhlR* mutant, which does not produce proteases. Clones with mutations in the PQS system displayed a bimodal phenotypic profile: 14 produced almost no proteases, while 7 clones had a similar or higher protease production level than the ancestral wild type.

Pyocyanin production is significantly reduced in all mutant categories as compared to the ancestral wild type (Fig. 2C, one-way ANOVA, F_5,191_ = 70.212, p < 0.001, all pairs tested with post-hoc Tukey HSD show P_adj_ < 0.001), but there are no significant differences between the mutant categories (all pairs tested with post-hoc Tukey HSD show P_adj_ > 0.500).

Rhamnolipid production was also significantly reduced in all mutant categories relative to the ancestral wild type (Fig. 2D, one-way ANOVA, F_5,191_ = 49.003, p < 0.001, all pairs tested with post-hoc Tukey HSD show P_adj_ < 0.001). But this time, we also observed significant differences in the production of rhamnolipids between the mutant categories (post-hoc Tukey HSD test, Las versus PQS and Las versus Las + PQS, P_adj_ < 0.001; Rhl versus Rhl + PQS, P_adj_ = 0.009). Clones with mutations in the PQS system stood out from the other categories because they showed enormous variability in rhamnolipid production spanning the entire continuum from zero to levels almost identical to the ancestral wild type.

Finally, when looking at the ability of these clones to form surface-attached biofilms, we found significant differences in biofilm production between the five mutant categories and the ancestral wild type (Fig. 2E, one-way ANOVA, F_5,191_ = 14.502, p < 0.001). While clones with Las mutations showed significantly reduced biofilm formation compared to the ancestral wild type (post-hoc Tukey HSD test, Las versus wild type, P_adj_ = 0.001; Las + PQS versus wild type, P_adj_ = 0.012), clones with Rhl and PQS mutations were on average not different from the ancestral wild type (post-hoc Tukey HSD test, Rhl versus wild type, P_adj_ = 0.859; PQS versus wild type, P_adj_ = 1.000). However, we observed again enormous variability among PQS mutants: while some mutants show extremely reduced biofilm formation, others invest considerably more into this trait compared to the ancestral wildtype.

The above findings suggest that mutations in the Las regulon spur broad-scale loss of function of QS traits, while mutations in the Rhl and PQS regulon modulate the QS-regulated trait expression patterns. To explore the apparent phenotypic segregation between mutant categories, we performed a principal component analysis (PCA) incorporating all five phenotypes into a single analysis (Fig. 2F). We found that the evolved clones significantly clustered based on the mutant categories (PERMANOVA; F_4,60_ = 28.167, p = 0.001). When focusing on the loadings of the first two principal components (PCs) (i.e., vectors in Fig. 2F, Table S3), we can identify two trade-offs among the QS-regulated traits. PC1 yields a trade-off between planktonic growth and biofilm formation as well as rhamnolipid production, meaning that evolved clones producing higher amounts of biofilm matrix components and rhamnolipids tend to grow less well in planktonic cultures. PC2 reveals a trade-off between protease and pyocyanin production, indicating that evolved clones that produce higher levels of proteases make lower levels of pyocyanin and vice versa. At the global level, we can conclude that modulation in the production of QS traits seems to be guided by trade-offs, meaning that maintaining or increasing the expression of one QS trait is associated with a proportional reduction of another QS trait. As QS modulations seem to be most marked among Rhl and PQS mutants, we focus more closely on these two QS systems in the next sections.

### Modulation of the Rhl regulon

All the three clones that have SNPs in the Rhl receptor, RhlR, have highly upregulated protease production, downregulated pyocyanin and rhamnolipid production, but retained wild type level formation of surface-attached biofilm. This phenotypic profile points towards QS regulon modulation, where the trait portfolio of these clones has changed. Here, we hypothesize that these phenotypic modulations should be reflected at the gene expression level. To test this, we used double fluorescent gene reporters to simultaneously measure transcriptional gene expression activity of *rhlR* and the receptor of the upstream Las system, *lasR,* in these three clones over a growth period of 24 hours in LB medium (Fig. 3A, Table S4). We found that mutations in *rhlR* significantly upregulated the expression of its own gene as compared to the wild type strain (one-way ANOVA: F_2,33_ = 54.950, p < 0.001; post-hoc Tukey HSD test, P_adj_ < 0.001). However, this upregulation did not occur in the two clones that had mutations in *pqsR* in addition to *rhlR* mutations (post-hoc Tukey HSD test, P_adj_ = 0.971). Curiously, we found that *lasR* expression was also significantly increased in clones with mutations in *rhlR,* and in one of the two clones with mutations in both *rhlR* and *pqsR* (one-way ANOVA, F_2,33_ = 22.554, p < 0.001). The expression trajectory of *lasR* in the four overexpressing clones follows a cyclical pattern with two successive expression peaks at hours 10 and 18. The second peak coincides with the expression peak observed in *rhlR*. Taken together, our results reveal that point mutations in *rhlR* can lead to highly increased gene expression levels of the QS receptors RhlR and LasR.

**Figure 3:**
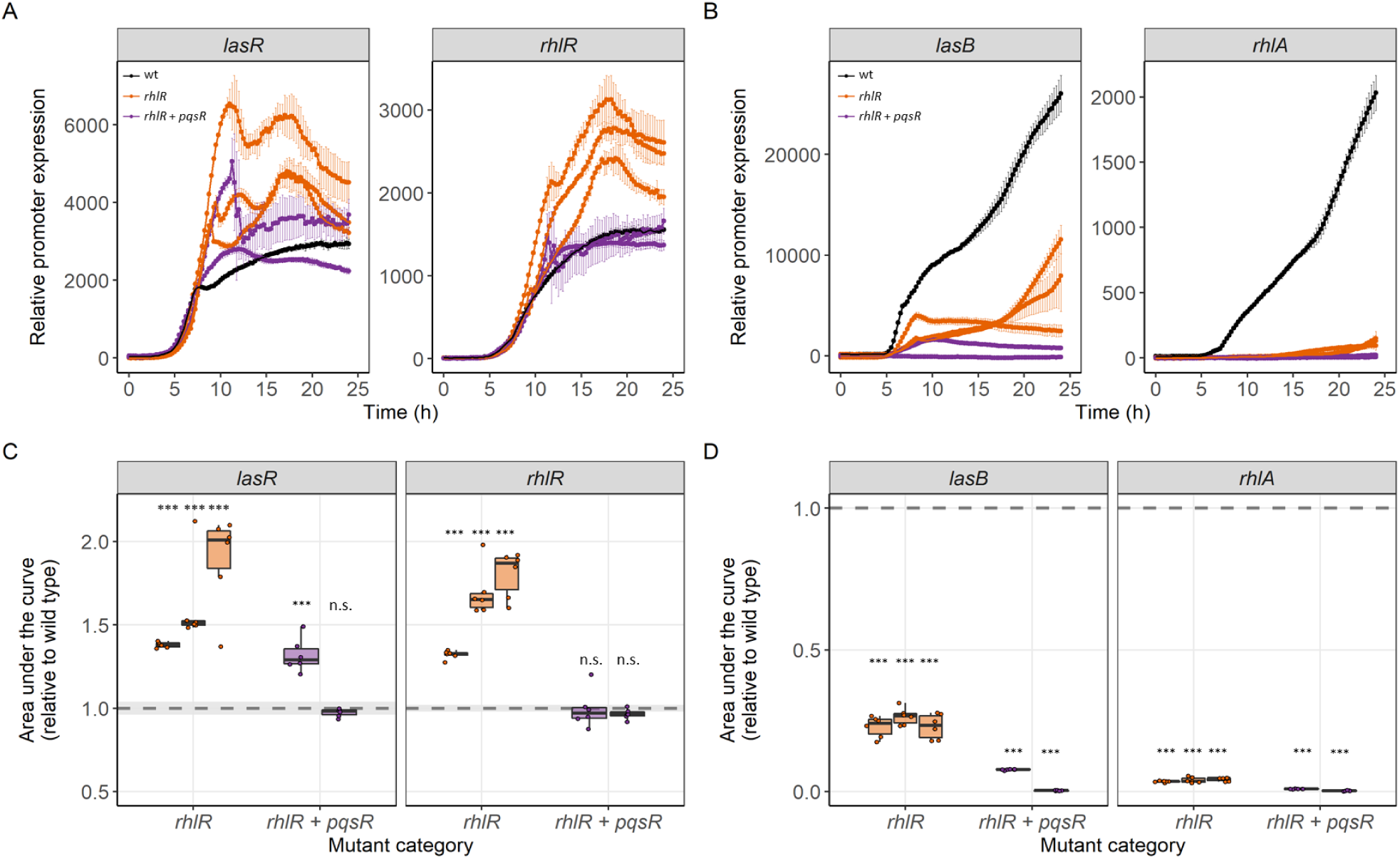
Mutations in *rhlR* upregulates expression of the Las and Rhl receptors. Gene expression trajectories of (A) Las (*lasR*) and Rhl (*rhlR*) receptors, and (B) Las-regulated protease (*lasB*) and Rhl-regulated rhamnolipid synthesis enzyme (*rhlA*) in clones with mutations in *rhlR* (orange) and *rhlR* + *pqsR* (purple) shown as means ± standard deviation. Gene expression in PAO1 wild type strain was used as reference control (black). Gene expression was measured as mCherry or GFP fluorescence and reported as fluorescence units, blank corrected by the background fluorescence of the wild type untagged strain. (C-D) Area under the gene expression trajectory curves in individual clones (represented by boxplots), relative to the gene expression in PAO1 wild type strain (dashed line at 1.0 ± standard deviation depicted as shaded area). Data stem from 6 independent replicates per clone. Asterisks indicate whether area under the curve is significantly different from the PAO1 wild type strain (based on post-hoc Tukey HSD: n.s. = not significant, *** p < 0.001).

Increased expression of QS receptors could lead to higher transcriptional regulator activity within the QS network, and further translate to an increased expression of the downstream QS genes (Fig. 3B). To test this hypothesis, we measured the expression of *lasB,* a protease regulated by both the Las and Rhl systems (Schuster, Urbanowski and Greenberg, 2004), and *rhlA*, part of the RhlAB rhamnolipid operon regulated by the Rhl system (Lequette and Greenberg, 2005). We found no support for our hypothesis, as the *lasB* expression level was reduced in all the five clones (one-way ANOVA, F_2,33_ = 1204.4, p < 0.001; post-hoc Tukey HSD test for all pairs, P_adj_ < 0.001). Similarly, we found strongly reduced expression of *rhlA* in all the five clones, with some of the expression levels being close to zero (one-way ANOVA, F_2,33_ = 8046.8, p < 0.001; post-hoc Tukey HSD test for all pairs, P_adj_ < 0.001). These findings show that the increased expression of the LasR and RhlR QS receptors do not translate into increased expression of two downstream regulated QS-traits, LasB protease and rhamnolipid synthesis enzymes. For *rhlA*, our gene expression results are compatible with the phenotypic data, as all mutants showed greatly reduced rhamnolipid production. For *lasB*, our gene expression results suggest that other proteases than LasB must be responsible for the observed high protease production at the phenotypic level.

### Modulation of the PQS regulon

Our phenotypic screening from Figure 2 revealed that mutations in the PQS system result in the most variable changes in the QS-regulated traits, with several clones showing an upregulation of QS traits. Here, we focus on the 21 clones that have mutations only in the PQS system to explore whether evolved QS phenotypes depend on the mutated gene within the PQS locus, and whether there are trade-offs, where the upregulation of one QS trait results in the downregulation of another one. Accordingly, we split the clones based on the mutated sites: PQS signal operon (*pqsABCD* and *pqsE*), PQS transcriptional regulator (*pqsR*) and double mutants and re-run our phenotypic analysis (Fig. 4A-E). We found that sample size was too small for most categories to reliably establish relationships between phenotypes and mutational patterns. However, when conducting a PCA with all clones we found that the evolved phenotypic profiles differed significantly between the mutation sites within the PQS regulon (PERMANOVA; F_3,20_ = 2.712, p = 0.031, Fig. 4F). We further observed two trade-offs among traits (Fig. 4F, Table S5). First, clones with higher levels of protease production and biofilm formation produced less pyocyanin (Fig. S1 A-B). Second, clones with higher levels of biofilm formation had lower growth in planktonic culture (Fig. S1 C).

**Figure 4.**
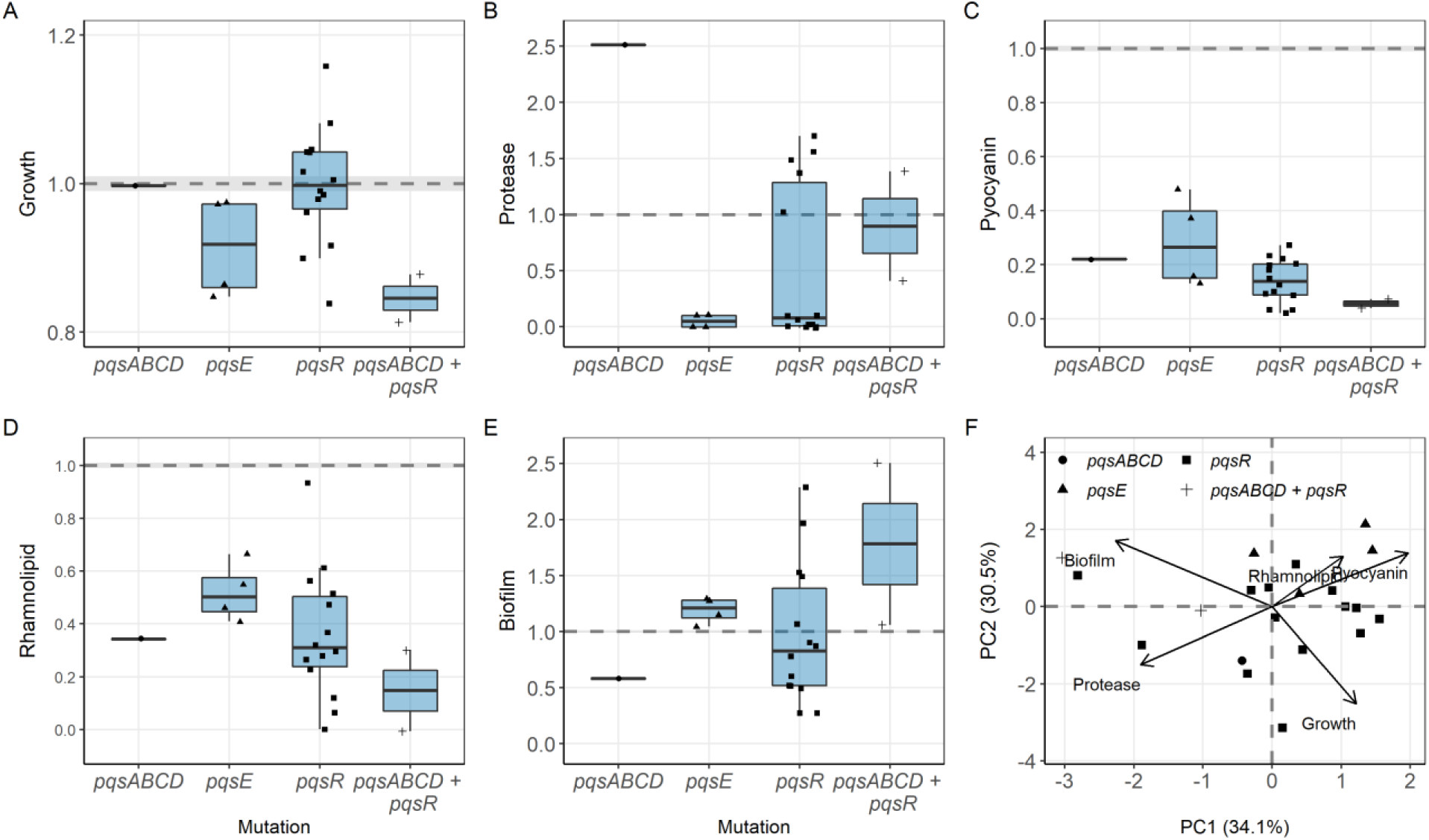
Phenotypes in PQS mutants and trade-offs in the production of QS-regulated traits. (A) Endpoint planktonic growth (OD at 600nm) in LB medium after 24 hours, and production of four QS-regulated traits: (B) protease, (C) pyocyanin, (D) rhamnolipid and (E) surface-attached biofilm. The values of measured phenotypes are expressed relative to the corresponding value of the ancestral PAO1 wild type strain (mean ± standard error indicated as dotted lines and shaded areas, respectively). (F) Principal component analysis (PCA) on the production of growth and four QS-regulated traits reveals significant clustering of mutant types and significant trade-offs (opposing vectors) between certain phenotypes. Each data point represents the average measure of at least three independent replicates per clone.

Finally, we had a closer look at the clones with mutations in *pqsR*, which represent the most frequent mutant type and show the highest variability for most phenotypes. Especially, the bimodal profile of protease production observed in Figure 2 is prevalent among the *pqsR* mutants (Fig. 4B). Here, we tested whether the divergent trajectories across *pqsR* mutants are linked to the location of the mutations (ligand- (LDB) versus DNA- (DBD) binding domain), or the type of mutations with regard to their deleterious effects (missense versus frameshift/deletions) (Table S6). However, we found that neither of these two factors can explain the bimodal protease production profiles (Fisher’s exact test: location of mutations, p = 1, type of mutations, p = 0.608).

## Discussion

As an important human pathogen, the evolution of *Pseudomonas aeruginosa* populations has been studied in a myriad of context, with extensive genetic adaptation being repeatedly observed in diverging environments such as human cystic fibrosis (CF) lungs, animal infection models, as well as in the natural environments and *in vitro* experimental evolutions (Rumbaugh *et al*., 2009; Jansen *et al*., 2015; Feltner *et al*., 2016; Granato *et al*., 2018; Groleau *et al*., 2021; Scribner *et al*., 2021; Smalley *et al*., 2022). The quorum sensing (QS) regulon, a global three-unit regulatory system that controls the expression of up to 10% of the genes in *P. aeruginosa*, many of which include virulence factors, is often among the most mutated pathways. It remains unclear why mutations in the QS regulon are consistently favored across different environments. QS mutants could arise and spread due to (I) disuse of the regulon, (II) cheating on the cooperative benefits of QS, or (III) modulation of one or several of the three systems. Here, we used a set of 61 experimentally evolved QS mutants (with mutations in the three systems: Las, Rhl and PQS) to examine these three scenarios. We found a clear distinction between the QS systems in how mutations affected the production of QS-regulated traits. While mutations in the Las system generally resulted in the loss of QS function (supporting scenarios I and II), mutations in the Rhl and PQS systems resulted in an altered trait production portfolio and regulatory network modulations (supporting scenario III).

The mutational and phenotypic patterns observed in the Las mutants strongly point towards loss of QS function as opposed to regulon modulation. Our findings are partially in contrast with previous studies showing that Las mutants can retain QS activity, partly through re-wiring the QS network (Feltner *et al*., 2016; Chen *et al*., 2019; Kostylev *et al*., 2019). We suggest that the difference in our findings is driven by the fact that we predominantly found large-scale deletions of the Las system, where the signal synthase (*lasI*), repressor (*rsaL*) and regulator (*lasR*) are deleted, therefore, leading to a loss of QS function. There is increasing evidence that large-scale deletions of the Las system are common in *in vitro* experimental evolution (O’Brien *et al*., 2017; Scribner *et al*., 2021; Tostado-Islas *et al*., 2021), but might have been overlooked in the past due to computational challenges of identifying them in draft genomes produced by short-read sequencing. Given that these large-scale deletions occurred many times independently, they must have an adaptive advantage. Similar large-scale deletions have previously been associated with both cheating and loss due to disuse in the process of medium adaptation (O’Brien *et al*., 2017; Scribner *et al*., 2021; Tostado-Islas *et al*., 2021). In our case, the QS mutants emerged from wild type populations that had been experimentally evolved in casamino acid medium, predominantly consisting of digested amino acids, an environment in which classic QS-regulated traits such as proteases and rhamnolipids are not needed. Thus, disuse is a plausible explanation for the selective spread of these mutants, even more so because the loss of QS fostered increased production of the siderophore pyoverdine, which was beneficial in the context of the initial study (Figueiredo, Wagner and Kümmerli, 2021). Taken together, whether loss of the Las system is caused by disuse or cheating is often context-dependent and spurred by the relative costs and benefits of the QS system in the respective environment.

We found surprisingly few mutants with point mutations in *lasR* compared to other studies (Smith *et al*., 2006; Jansen *et al*., 2015; Granato *et al*., 2018; Chen *et al*., 2019). Such mutants are likely to produce structural variants of the LasR master regulator, which could form the basis of QS regulon modulation. However, we found little evidence for this, as our clones with point mutations in *lasR* showed similar trait portfolios as the large-scale Las deletion mutants with greatly reduced or completely abolished production of QS-regulated traits (Fig. 2). Thus, they are most likely loss-of-function mutants that spread because of either disuse or cheating. In the latter case, they can be considered as signal-blind (*lasR*) mutants that save the costs of cooperating while exploiting the cooperative signaling and exoproduct production by QS-wild type individuals (West *et al*., 2006; Diggle *et al*., 2007; Wilder, Diggle and Schuster, 2011).

In contrast, we found evidence for QS regulon modulation in the Rhl mutants. These mutants arose at a much lower frequency than the Las mutants, similar to previous studies (Bjarnsholt *et al*., 2010; Ahmed *et al*., 2021). Both the phenotypic profiles and gene expression analyses revealed that there is no complete loss of QS function, but rather a change in the expression of the QS-regulated trait portfolio. At the phenotypic level, the regulon modulation is characterized by upregulating protease production and downregulating pyocyanin and rhamnolipid production, while retaining the ability to form surface-attached biofilms at levels similar to the ancestral wild type. This points towards possible decoupling of certain elements from the QS regulon. A straightforward explanation would be that mutations in *rhlR* abolish the production of traits that are directly controlled by the Rhl system like phenazines and rhamnolipids, whilst maintaining the traits that are predominantly under the control of the hierarchically superior Las system like proteases. However, our data speak against this explanation as we found largely increased protease production and significantly increased *rhlR* and *lasR* gene expression (Fig. 3), suggesting that the QS regulon is modulated in a more complicated way. One alternative explanation is that mutations in *rhlR* give rise to RhlR receptor variants that, upon binding to the signal, show altered transcriptional factor affinities to promoter binding sites. In other words, mutated RhlR variants could trigger increased expression of certain traits (including its own expression), while others are downregulated. Such regulon modulations seem to take non-linear paths as indicated by our data showing that increased *rhlR* expression is associated with increased *lasR* expression, but decreased expression of *lasB*, which is directly controlled by LasR. This finding matches our observation in the wild type *P. aeruginosa*, where we found high levels of *lasR* expression in a fraction of clonal cells is associated with low *lasB* expression (Jayakumar *et al*., 2021). However, reduced expression levels of *lasB* do not correspond with our observation that these Rhl mutants have higher protease production (Fig. 2), suggesting that other important QS-regulated proteases, such as LasA and AprA (Gambello, Kaye and Iglewski, 1993; Coin *et al*., 1997) might be upregulated instead of LasB. Taken together, while we found evidence for mutations in *rhlR* leading to QS modulation, further research is required to unravel its complex causes and consequences.

Our analysis on PQS mutants reveal that these mutants are quite common (similar in frequency to the Las mutants) and show clear evidence for QS regulon modulation rather than loss of QS function. Unlike the Rhl mutants that all had similar changes in their trait production portfolio, the PQS mutants instead show a heterogeneous profile, with phenotypic trade-offs between some of the QS-regulated traits. For example, while a subset of PQS mutants produces higher amounts of surface-attached biofilms, others grow better in planktonic cultures or produce higher amounts of pyocyanin. At the same time, the PQS mutants segregate along a continuum from low protease but high pyocyanin production to high protease but low pyocyanin production. These phenotypic trade-offs open two interesting possibilities, namely when mutants with opposing phenotypes occur in the same population. First, the different mutations in the PQS regulon could spur diversification, whereby the various mutants follow successful strategies in different ecological niches. Second, mutants with diverging phenotypes may each specialize in the production of a set of QS-regulated traits and share these traits with the other specialists at the group level. While our sample size is too small to draw strong conclusions, we indeed found cases where such phenotypically divergent clones occur in the same population (Fig. S1). As for the Las and the Rhl systems, we observed that the large majority of mutations occurred in *pqsR,* that encodes for the regulator of the PQS system. Therefore, we propose that mutations in this gene results in PqsR receptor variants that in combination with the signal show differential affinities as transcription factor, leading to the increased expression of certain traits and the downregulation of other traits. But interestingly, and different from the Las and Rhl systems, we also found mutations in other genes of the PQS regulon (*pqsABCD* and *pqsE*), which seem to contribute to the diversification observed at the phenotypic level (Fig. 4). As with the Rhl mutants, modulation of the PQS network is indeed intricate and further genetic work is required to elucidate the exact regulatory trajectories that drive the altered trait expression profile among the PQS mutants.

Finally, we also had a low number of clones with mutations in two QS systems (Las + PQS, n = 2; Rhl + PQS, n = 2). Phenotypes of the Las + PQS mutants seem to be dominated by mutations in the Las system, leading to loss of QS function. Similarly, the phenotypes of Rhl + PQS mutants point more towards loss of QS function rather than regulon modulation. For example, while single Rhl or PQS mutants produce rhamnolipids and can form surface-attached biofilms, albeit at varying levels, the double mutants show abolished phenotypes. The damping of QS modulation in double mutants is also observed at the gene expression level, where the Rhl + PQS mutants did not show an increased *rhlR* expression as observed for the Rhl single mutants (Fig. 3). This indicates that an active PQS system is required for the regulon modulation to work in Rhl mutants. Our results are in line with previous studies reporting that mutation in the PQS regulon can result in a partial loss of Rhl activity, most likely through the disruption of an alternative signaling molecule that is recognized by RhlR (Mukherjee *et al*., 2018; Kostylev *et al*., 2019).

In conclusion, our results reinforce the view that QS is under selection not only in infections but also in *in vitro* experimental evolution. We show that mutational patterns and resulting phenotypes are complex. While mutations in the Las system typically are associated with loss of QS function, we find that mutations in the Rhl and PQS systems lead to regulon modifications. The idea that QS network can evolve and be rewired to match prevailing conditions in the laboratory and the host has only recently emerged (Jansen *et al*., 2015; Feltner *et al*., 2016; Oshri *et al*., 2018; Chen *et al*., 2019; Kostylev *et al*., 2019). Here, we lend support to this hypothesis. The next goal is to understand how QS regulon modulation affects the plasticity and flexibility in coordinating the social phenotypes in *P. aeruginosa* and what the consequences of QS modulation are for bacterial fitness and virulence in the host. Our study yields first indications that modulation may drive strain diversification and adaptation to different ecological niches and may perhaps also foster mutualist interactions between emerging strains (Rezzoagli, Granato and Kümmerli, 2020). Future research should investigate these aspects and extend the search for mutations to accessory regulatory elements of the QS network, such as VqsM, AlgR and Vfr (Morici *et al*., 2007; Folkesson *et al*., 2012; Liang *et al*., 2014), which may further contribute to regulon modulation.

## Acknowledgements

We thank Richard Allen for help with statistical analysis. This work was funded by the European Research Council (ERC) under the European Union’s Horizon 2020 research and innovation program (grant agreement no. 681295), the Swiss National Science Foundation (grant no. 31003A_182499), and a grant from the University Research Priority Program “Evolution in Action”.

## Author contribution

P.J., A.R.T.F. and R.K. designed the study; P.J. performed the experiments and analysed the data; P.J., A.R.T.F. and R.K. interpreted the data and wrote the paper.

## SUPPLEMENTARY MATERIAL

**Supplementary Figure 1:**
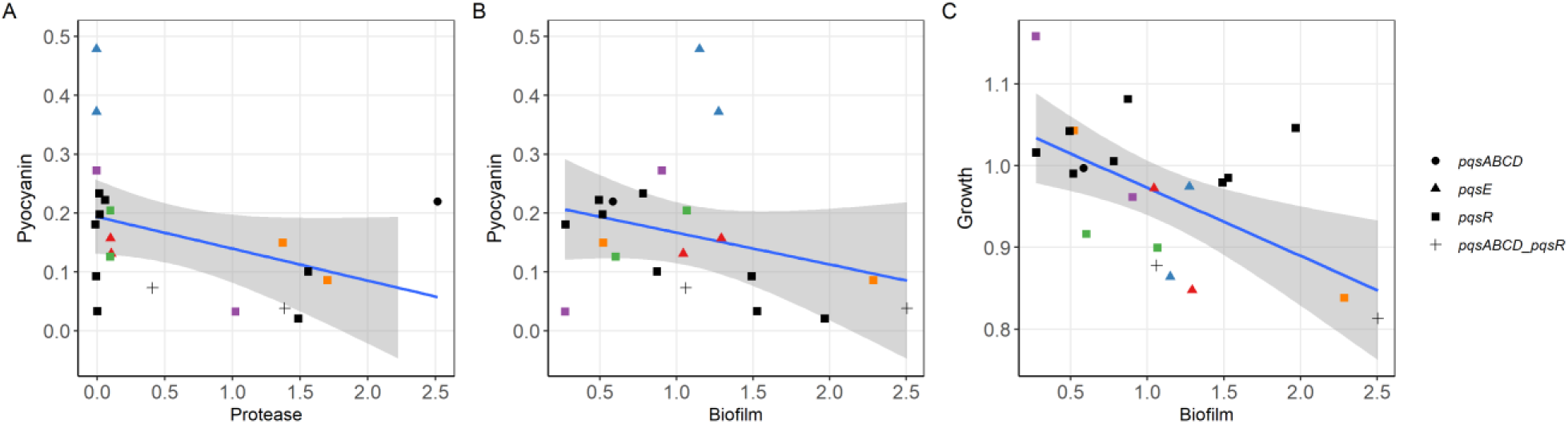
Trade-offs between traits in PQS mutants. Associations are shown for (A) production of pyocyanin versus protease, (B) production of pyocyanin versus ability to form surface-attached biofilms, and (C) planktonic growth versus ability to form surface-attached biofilms. Coloured symbols represent clones originating from the same evolved population. All other clones are coloured black. Each data point represents the average measure of at least three independent replicates per clone. Grey shaded region represents 95% confidence interval.

**Table S1.**
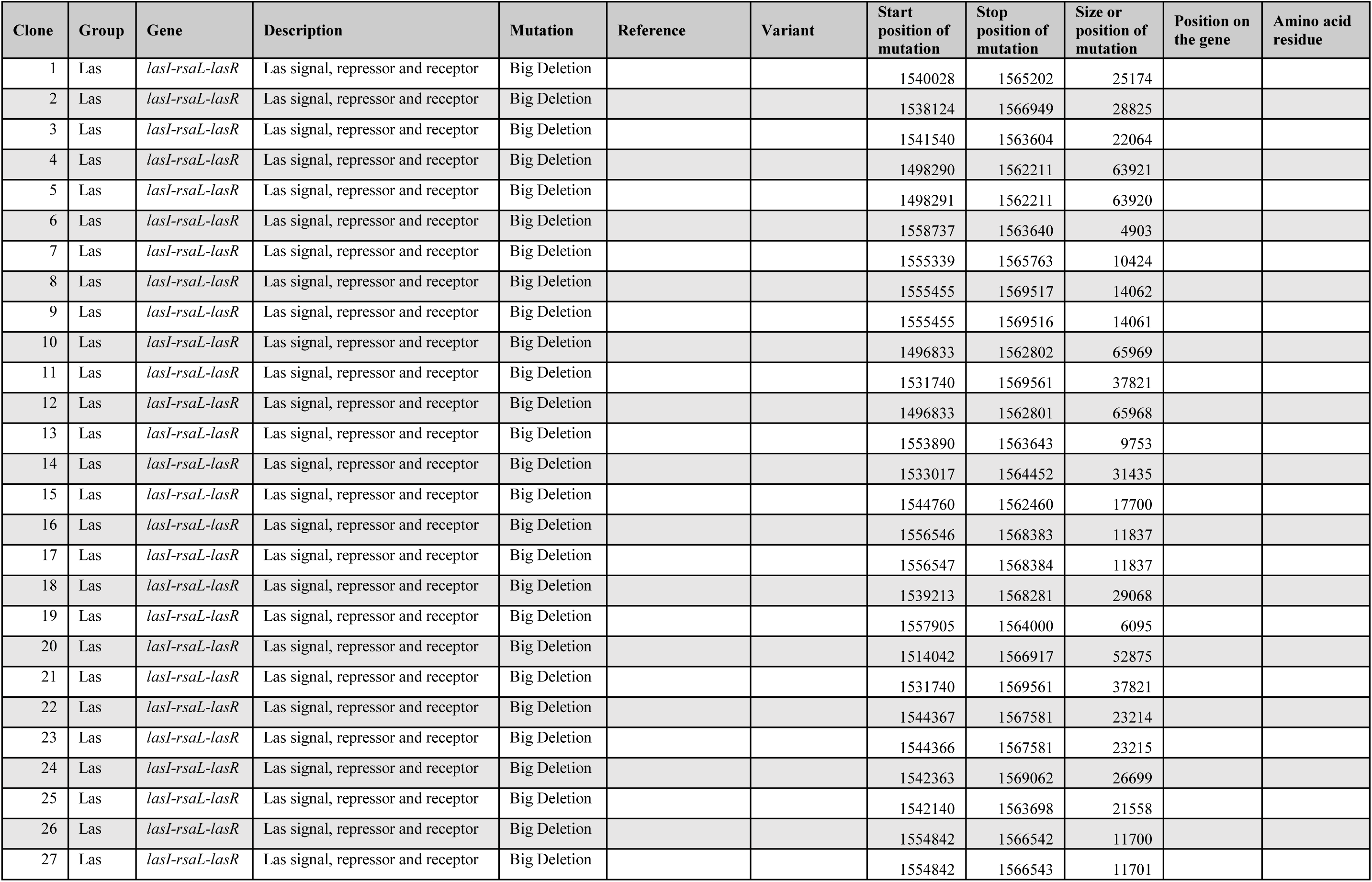
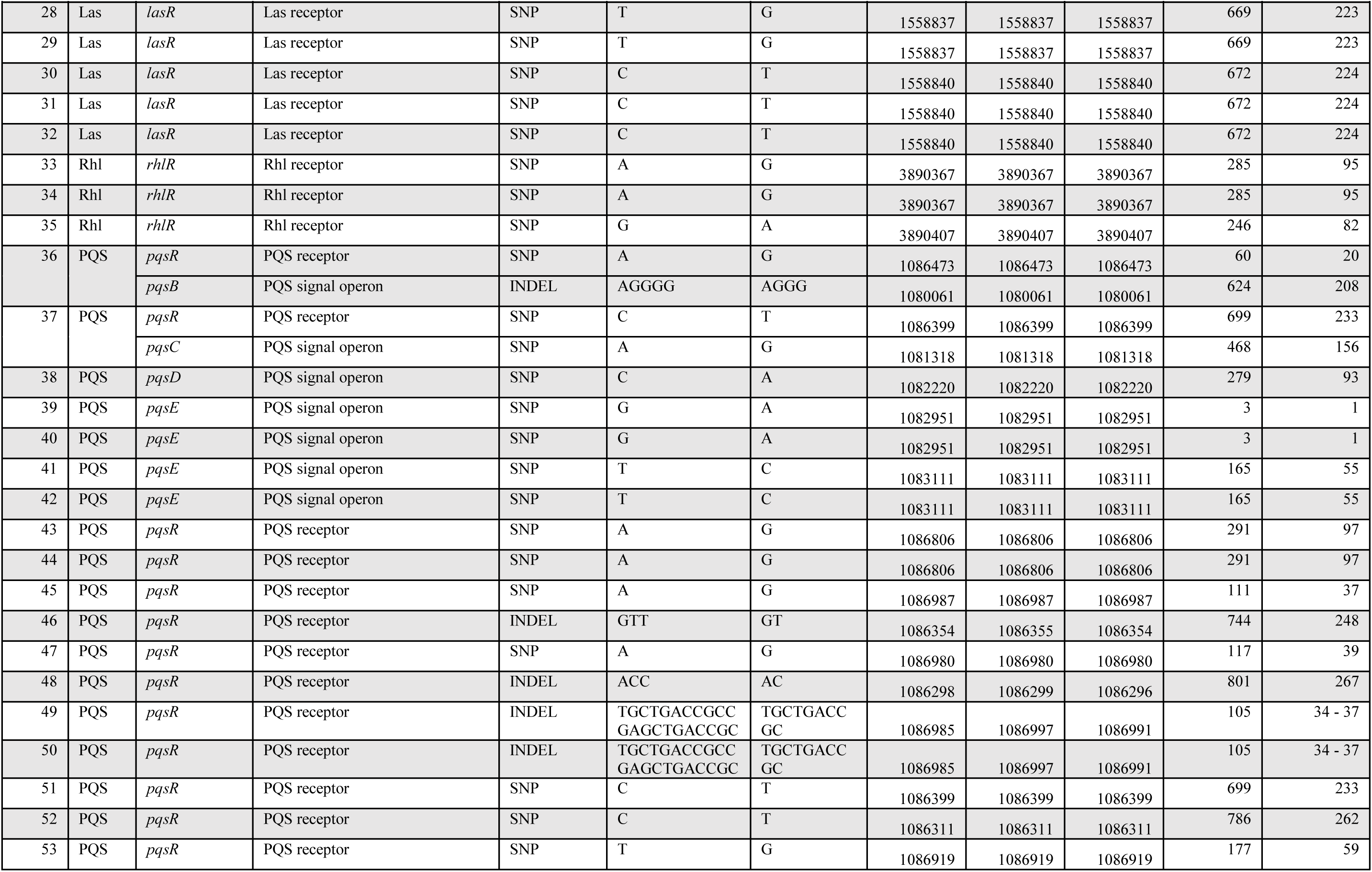
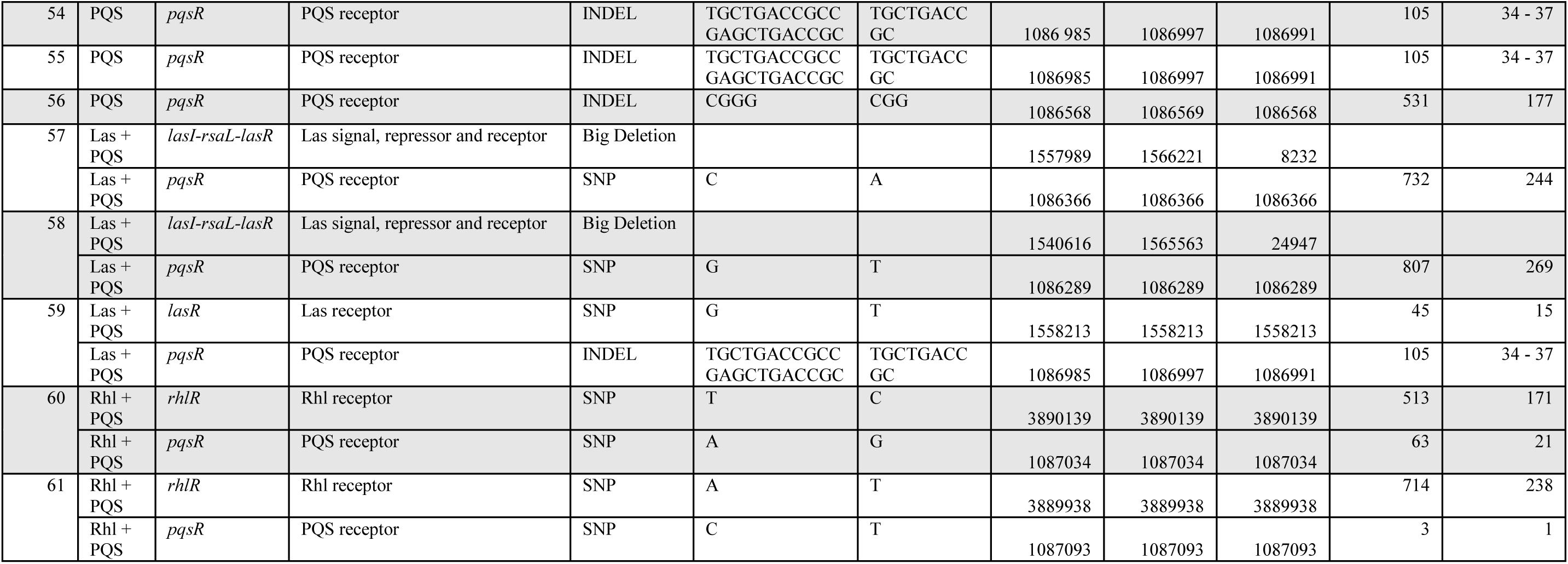
List of experimentally evolved *Pseudomonas aeruginosa* PAO1 strains

**Table S2.** List of defined bacterial strains and plasmids used for strain construction and for control experiments.

| Strains | Description | Source or reference |
| --- | --- | --- |
| ***E. coli*** |  |  |
| CC118 λpir | $\Delta$ (*ara*, *leu*)<sub>7697</sub> *araD*139 Δ*lacX*74 *galE galK phoA*20 *thi-1 rpsE rpoB* (Rf<sup>R</sup>) *argE(am) recA1* λpir<sup>+</sup> | De Lorenzo *et al.*, 1990 |
| ***Pseudomonas aeruginosa*** |  |  |
| PAO1 wild type | Wild type strain (ATCC 15692) | This laboratory |
| PAO1 Δ*lasR* | Deficient in the receptor of Las system | This laboratory |
| PAO1 Δ*rhlR* | Deficient in the receptor of Rhl system | This laboratory |
| PAO1 Δ*lasR* Δ*rhlR* | Deficient in the receptor of both Las and Rhl systems | This laboratory |
| **Plasmids** |  |  |
| pUX-BF13 | Helper plasmid to provide Tn7 transposase proteins for reporter construct integration at the *attTn7* site in *P. aeruginosa* | Bao *et al.*, 1991 |
| pDR05 | pUC18-mini-Tn7-Gm with *lasR-GFP* and *rhlR-mCherry* | Jayakumar *et al.*, 2021 |
| pDR06 | pUC18-mini-Tn7-Gm with *lasB-GFP* and *rhlA-mCherry* | Jayakumar *et al.*, 2021 |

**Table S3:** Loadings of phenotypic variables onto the principal components (PCs).

|  | PC1 | PC2 | PC3 | PC4 | PC5 |
| --- | --- | --- | --- | --- | --- |
| Growth | 0.3557 | -0.3695 | 0.7588 | 0.2855 | 0.2823 |
| Protease | -0.2117 | 0.5752 | 0.5797 | -0.5369 | 0.0043 |
| Pyocyanin | -0.2236 | -0.6773 | -0.0374 | -0.6764 | 0.1800 |
| Rhamnolipid | -0.5975 | -0.2593 | 0.293 | 0.269 | -0.6462 |
| Biofilm | -0.6494 | 0.0819 | -0.0301 | 0.3168 | 0.6858 |
| Explained variance (%) | 32.9 | 24.6 | 18.3 | 14.7 | 9.5 |

**Table S4:** List of fluorescent gene reporter strains

| Strains | Description | Source or reference |
| --- | --- | --- |
| PAO1 wild type<br>*promoterless-GFP and mCherry* | Non-fluorescent wild type strain with an empty promoter site fused to GFP and mCherry | Jayakumar *et al.*, 2021 |
| **Evolved *rhlR* mutants** |  |  |
| Clone 33-*lasR-gfp-rhlR-mCherry* | Transcriptional fusion of *lasR-GFP* and *rhlR-mCherry* from pDR05 | This study |
| Clone 33-*lasB-gfp-rhlA-mCherry* | Transcriptional fusion *lasB-GFP* and *rhlA-mCherry* from pDR06 | This study |
| Clone 34-*lasR-gfp-rhlR-mCherry* | Transcriptional fusion of *lasR-GFP* and *rhlR-mCherry* from pDR05 | This study |
| Clone 34-*lasB-gfp-rhlA-mCherry* | Transcriptional fusion *lasB-GFP* and *rhlA-mCherry* from pDR06 | This study |
| Clone 35-*lasR-gfp-rhlR-mCherry* | Transcriptional fusion of *lasR-GFP* and *rhlR-mCherry* from pDR05 | This study |
| Clone 35-*lasB-gfp-rhlA-mCherry* | Transcriptional fusion *lasB-GFP* and *rhlA-mCherry* from pDR06 | This study |
| **Evolved *rhlR* mutants containing mutations in *pqsR*** |  |  |
| Clone 62-*lasR-gfp-rhlR-mCherry* | Transcriptional fusion of *lasR-GFP* and *rhlR-mCherry* from pDR05 | This study |
| Clone 62-*lasB-gfp-rhlA-mCherry* | Transcriptional fusion *lasB-GFP* and *rhlA-mCherry* from pDR06 | This study |
| Clone 63-*lasR-gfp-rhlR-mCherry* | Transcriptional fusion of *lasR-GFP* and *rhlR-mCherry* from pDR05 | This study |
| Clone 63-*lasB-gfp-rhlA-mCherry* | Transcriptional fusion *lasB-GFP* and *rhlA-mCherry* from pDR06 | This study |

**Table S5:** Loadings of phenotypic variables of *pqs* mutants onto the principal components (PCs).

|  | PC1 | PC2 | PC3 | PC4 | PC5 |
| --- | --- | --- | --- | --- | --- |
| Growth | 0.3119 | -0.6463 | 0.3768 | 0.0073 | 0.5856 |
| Protease | -0.4887 | -0.3860 | 0.1128 | -0.7395 | -0.2291 |
| Pyocyanin | 0.5054 | 0.3571 | -0.2834 | -0.6613 | 0.3153 |
| Rhamnolipid | 0.2635 | 0.3355 | 0.8406 | -0.1252 | -0.3094 |
| Biofilm | -0.5822 | 0.4396 | 0.2416 | -0.0061 | 0.6398 |
| Explained variance (%) | 34.1 | 30.5 | 19.4 | 12.6 | 3.5 |

**Table S6:** Clones with mutations in the PQS regulator (*pqsR*)

| Clone number | Protease production | Protein Domain | Mutation type |
| --- | --- | --- | --- |
| 36 | Low | DBD | Frameshift |
| 43 | Low | LBD | Missense |
| 44 | Low | LBD | Missense |
| 45 | Low | DBD | Missense |
| 47 | Low | DBD | Missense |
| 48 | Low | LBD | Frameshift |
| 49 | Low | DBD | Conservative in-frame deletion |
| 52 | Low | LBD | Missense |
| 53 | Low | DBD | Missense |
| 56 | Low | LBD | Frameshift |
| 37 | High | DBD | Missense |
| 46 | High | DBD | Frameshift |
| 50 | High | LBD | Conservative in-frame deletion |
| 51 | High | DBD | Missense |
| 54 | High | LBD | Conservative in-frame deletion |
| 55 | High | LBD | Conservative in-frame deletion |

\*LBD: Ligand-binding domain; DBD: DNA-binding domain

## References

Ahmed, S. A. K. S. et al. (2021) ‘*Pseudomonas aeruginosa* PA80 is a cystic fibrosis isolate deficient in RhlRI quorum sensing’, Scientific Reports, 11(1), p. 5729. doi: 10.1038/s41598-021-85100-0.

Bjarnsholt, T. et al. (2010) ‘Quorum sensing and virulence of *Pseudomonas aeruginosa* during lung infection of cystic fibrosis patients’, PLoS ONE. PLoS One, 5(4). doi: 10.1371/journal.pone.0010115.

Bodour, A. A. and Miller-Maier, R. M. (1998) ‘Application of a modified drop-collapse technique for surfactant quantitation and screening of biosurfactant-producing microorganisms’, Journal of Microbiological Methods. Elsevier, 32(3), pp. 273–280. doi: 10.1016/S0167-7012(98)00031-1.

Chen, R. et al. (2019) ‘Social cheating in a *Pseudomonas aeruginosa* quorum-sensing variant’, Proceedings of the National Academy of Sciences of the United States of America, 116(14), pp. 7021–7026. doi: 10.1073/pnas.1819801116.

Coin, D. et al. (1997) ‘LasA, alkaline protease and elastase in clinical strains of *Pseudomonas aeruginosa*: Quantification by immunochemical methods’, FEMS Immunology and Medical Microbiology, 18(3), pp. 175–184. doi: 10.1016/S0928-8244(97)00037-0.

Cruz, R. L. et al. (2020) ‘RhlR-regulated acyl-homoserine lactone quorum sensing in a cystic fibrosis isolate of *Pseudomonas aeruginosa*’, mBio, 11(2). doi: 10.1128/mBio.00532-20.

D’Argenio, D. A. et al. (2007) ‘Growth phenotypes of *Pseudomonas aeruginosa lasR* mutants adapted to the airways of cystic fibrosis patients’, Molecular Microbiology, 64(2), pp. 512–533. doi: 10.1111/J.1365-2958.2007.05678.X.

Damkiær, S. et al. (2013) ‘Evolutionary remodeling of global regulatory networks during long-term bacterial adaptation to human hosts’, Proceedings of the National Academy of Sciences of the United States of America. Proc Natl Acad Sci U S A, 110(19), pp. 7766–7771. doi: 10.1073/pnas.1221466110.

Dettman, J. R. et al. (2013) ‘Evolutionary genomics of epidemic and nonepidemic strains of *Pseudomonas aeruginosa*’, Proceedings of the National Academy of Sciences of the United States of America, 110(52), pp. 21065–21070. doi: 10.1073/pnas.1307862110.

Diggle, S. P. et al. (2007) ‘Cooperation and conflict in quorum-sensing bacterial populations’, Nature, 450(7168), pp. 411–414. doi: 10.1038/nature06279.

Feltner, J. B. et al. (2016) ‘LasR variant cystic fibrosis isolates reveal an adaptable quorum-sensing hierarchy in *Pseudomonas aeruginosa*’, mBio. American Society for Microbiology, 7(5), pp. e01513–16. doi: 10.1128/mBio.01513-16.

Figueiredo, A. R. T., Wagner, A. and Kümmerli, R. (2021) ‘Ecology drives the evolution of diverse social strategies in *Pseudomonas aeruginosa*’, Molecular Ecology. John Wiley & Sons, Ltd, 30(20), pp. 5214–5228. doi: 10.1111/mec.16119.

Folkesson, A. et al. (2012) ‘Adaptation of *Pseudomonas aeruginosa* to the cystic fibrosis airway: an evolutionary perspective’, Nature Reviews Microbiology 2012 10:12. Nature Publishing Group, 10(12), pp. 841–851. doi: 10.1038/nrmicro2907.

Gambello, M. J., Kaye, S. and Iglewski, B. H. (1993) ‘LasR of *Pseudomonas aeruginosa* is a transcriptional activator of the alkaline protease gene (apr) and an enhancer of exotoxin A expression’, Infection and Immunity, 61(4), pp. 1180–1184. doi: 10.1128/iai.61.4.1180-1184.1993.

Granato, E. T. et al. (2018) ‘Low spatial structure and selection against secreted virulence factors attenuates pathogenicity in *Pseudomonas aeruginosa*’, ISME Journal. Springer US, 12(12), pp. 2907–2918. doi: 10.1038/s41396-018-0231-9.

Groleau, M. C. et al. (2021) ‘*Pseudomonas aeruginosa* isolates defective in function of the LasR quorum sensing regulator are frequent in diverse environmental niches’, Environmental Microbiology. John Wiley & Sons, Ltd. doi: 10.1111/1462-2920.15745.

Hammond, J. H. et al. (2016) ‘ Environmentally Endemic *Pseudomonas aeruginosa* Strains with Mutations in *lasR* Are Associated with Increased Disease Severity in Corneal Ulcers ’, mSphere. mSphere, 1(5). doi: 10.1128/msphere.00140-16.

Jansen, G. et al. (2015) ‘Evolutionary transition from pathogenicity to commensalism: Global regulator mutations mediate fitness gains through virulence attenuation’, Molecular Biology and Evolution. Mol Biol Evol, 32(11), pp. 2883–2896. doi: 10.1093/molbev/msv160.

Jayakumar, P. et al. (2021) ‘*Pseudomonas aeruginosa* reaches collective decisions via transient segregation of quorum sensing activities across cells’, bioRxiv, p. 2021.03.22.436499. doi: 10.1101/2021.03.22.436499.

Koch, C. and H o iby, N. (1993) ‘Pathogenesis of cystic fibrosis’, The Lancet. Elsevier, 341(8852), pp. 1065–1069. doi: 10.1016/0140-6736(93)92422-P.

Köhler, T., Buckling, A. and Van Delden, C. (2009) ‘Cooperation and virulence of clinical *Pseudomonas aeruginosa* populations’, Proceedings of the National Academy of Sciences. National Academy of Sciences, 106(15), pp. 6339–6344. doi: 10.1073/PNAS.0811741106.

Kostylev, M. et al. (2019) ‘Evolution of the *Pseudomonas aeruginosa* quorum-sensing hierarchy’, Proceedings of the National Academy of Sciences of the United States of America, 116(14), pp. 7027–7032. doi: 10.1073/pnas.1819796116.

Kramer, J., López Carrasco, M. Á. and Kümmerli, R. (2020) ‘Positive linkage between bacterial social traits reveals that homogeneous rather than specialised behavioral repertoires prevail in natural Pseudomonas communities’, FEMS Microbiology Ecology. Oxford University Press, 96(1). doi: 10.1093/femsec/fiz185.

Lee, J. and Zhang, L. (2015) ‘The hierarchy quorum sensing network in *Pseudomonas aeruginosa*’, Protein and Cell, 6(1), pp. 26–41. doi: 10.1007/s13238-014-0100-x.

Lequette, Y. and Greenberg, E. P. (2005) ‘Timing and localization of rhamnolipid synthesis gene expression in *Pseudomonas aeruginosa* biofilms.’, Journal of bacteriology. American Society for Microbiology (ASM), 187(1), pp. 37–44. doi: 10.1128/JB.187.1.37-44.2005.

Lesprit, P. et al. (2003) ‘Role of the quorum-sensing system in experimental pneumonia due to *Pseudomonas aeruginosa* in rats’, American journal of respiratory and critical care medicine. Am J Respir Crit Care Med, 167(11), pp. 1478–1482. doi: 10.1164/RCCM.200207-736BC.

Liang, H. et al. (2014) ‘Molecular mechanisms of master regulator VqsM mediating quorum-sensing and antibiotic resistance in *Pseudomonas aeruginosa*’, Nucleic Acids Research. Oxford University Press, 42(16), p. 10307. doi: 10.1093/NAR/GKU586.

Marvig, R. L. et al. (2015) ‘Convergent evolution and adaptation of *Pseudomonas aeruginosa* within patients with cystic fibrosis’, Nature Genetics. Nature Publishing Group, 47(1), pp. 57–64. doi: 10.1038/ng.3148.

Morici, L. A. et al. (2007) ‘*Pseudomonas aeruginosa* AlgR Represses the Rhl Quorum-Sensing System in a Biofilm-Specific Manner’, Journal of Bacteriology. American Society for Microbiology (ASM), 189(21), p. 7752. doi: 10.1128/JB.01797-06.

Mukherjee, S. et al. (2018) ‘The PqsE and RhlR proteins are an autoinducer synthase–receptor pair that control virulence and biofilm development in *Pseudomonas aeruginosa*’, Proceedings of the National Academy of Sciences of the United States of America. National Academy of Sciences, 115(40), pp. E9411–E9418. doi: 10.1073/pnas.1814023115.

Nadal Jimenez, P., et al. (2012) ‘The Multiple Signaling Systems Regulating Virulence in *Pseudomonas aeruginosa*’, Microbiology and Molecular Biology Reviews, 76(1), pp. 46–65. doi: 10.1128/mmbr.05007-11.

O’Brien, S. et al. (2017) ‘Adaptation to public goods cheats in *Pseudomonas aeruginosa*’, Proceedings of the Royal Society B: Biological Sciences. Royal Society Publishing, 284(1859). doi: 10.1098/rspb.2017.1089.

Oksanen, J. et al. (2020) ‘Package “vegan” Title Community Ecology Package Version 2.5-7’, R, 2.5(7), pp. 1–286. Available at: https://github.com/vegandevs/vegan/issues%5Cnhttps://cran.r-project.org,%5Cnhttps://github.com/vegandevs/vegan (Accessed: 24 January 2022).

Oshri, R. D. et al. (2018) ‘Selection for increased quorum-sensing cooperation in *Pseudomonas aeruginosa* through the shut-down of a drug resistance pump’, The ISME Journal 2018 12:10. Nature Publishing Group, 12(10), pp. 2458–2469. doi: 10.1038/s41396-018-0205-y.

Parkins, M. D., Somayaji, R. and Waters, V. J. (2018) ‘Epidemiology, biology, and impact of clonal *Pseudomonas aeruginosa* infections in cystic fibrosis’, Clinical Microbiology Reviews. American Society for Microbiology (ASM), 31(4). doi: 10.1128/CMR.00019-18.

Pearson, J. P. et al. (2000) ‘*Pseudomonas aeruginosa* cell-to-cell signaling is required for virulence in a model of acute pulmonary infection’, Infection and immunity. Infect Immun, 68(7), pp. 4331–4334. doi: 10.1128/IAI.68.7.4331-4334.2000.

Preston, M. J. et al. (1997) ‘Contribution of proteases and LasR to the virulence of *Pseudomonas aeruginosa* during corneal infections’, Infection and immunity. Infect Immun, 65(8), pp. 3086–3090. doi: 10.1128/IAI.65.8.3086-3090.1997.

Rezzoagli, C., Granato, E. T. and Kümmerli, R. (2020) ‘Harnessing bacterial interactions to manage infections: a review on the opportunistic pathogen *Pseudomonas aeruginosa* as a case example’, Journal of Medical Microbiology, 69, pp. 147–161. doi: 10.1099/jmm.0.001134.

Rumbaugh, K. P. et al. (1999) ‘Contribution of quorum sensing to the virulence of *Pseudomonas aeruginosa* in burn wound infections’, Infection and immunity. Infect Immun, 67(11), pp. 5854–5862. doi: 10.1128/IAI.67.11.5854-5862.1999.

Rumbaugh, K. P. et al. (2009) ‘Quorum Sensing and the Social Evolution of Bacterial Virulence’, Current Biology. Curr Biol, 19(4), pp. 341–345. doi: 10.1016/j.cub.2009.01.050.

Savoia, D. and Zucca, M. (2007) ‘Clinical and environmental Burkholderia strains: biofilm production and intracellular survival’, Current microbiology. Curr Microbiol, 54(6), pp. 440–444. doi: 10.1007/S00284-006-0601-9.

Schaber, J. A. et al. (2004) ‘Analysis of quorum sensing-deficient clinical isolates of *Pseudomonas aeruginosa*’, Journal of medical microbiology. J Med Microbiol, 53(Pt 9), pp. 841–853. doi: 10.1099/JMM.0.45617-0.

Schuster, M., Urbanowski, M. L. and Greenberg, E. P. (2004) Promoter specificity in Pseudomonas aeruginosa quorum sensing revealed by DNA binding of purified LasR, Proceedings of the National Academy of Sciences of the United States of America. doi: 10.1073/pnas.0407229101.

Scribner, M. R. et al. (2021) ‘The nutritional environment is sufficient to select coexisting biofilm and quorum-sensing mutants of *Pseudomonas aeruginosa*’, bioRxiv, p. 2021.09.01.458652. doi: 10.1101/2021.09.01.458652.

Smalley, N. E. et al. (2022) ‘Evolution of the Quorum Sensing Regulon in Cooperating Populations of *Pseudomonas aeruginosa*’, mBio. Edited by M. Whiteley. American Society for Microbiology1752 N St., N.W., Washington, DC, 13(1). doi: 10.1128/MBIO.00161-22.

Smith, E. E. et al. (2006) ‘Genetic adaptation by *Pseudomonas aeruginosa* to the airways of cystic fibrosis patients’, Proceedings of the National Academy of Sciences. National Academy of Sciences, 103(22), pp. 8487–8492. doi: 10.1073/PNAS.0602138103.

Tostado-Islas, O. et al. (2021) ‘Iron limitation by transferrin promotes simultaneous cheating of pyoverdine and exoprotease in *Pseudomonas aeruginosa*’, The ISME Journal 2021 15:8. Nature Publishing Group, 15(8), pp. 2379–2389. doi: 10.1038/s41396-021-00938-6.

Vanderwoude, J. et al. (2020) ‘ The evolution of virulence in *Pseudomonas aeruginosa* during chronic wound infection ’, Proceedings of the Royal Society B: Biological Sciences, 287(1937), p. 20202272. doi: 10.1098/rspb.2020.2272.

West, S. A. et al. (2006) ‘Social evolution theory for microorganisms’, Nature Reviews Microbiology. Nature Publishing Group, 4(8), pp. 597–607. doi: 10.1038/nrmicro1461.

Wilder, C. N., Diggle, S. P. and Schuster, M. (2011) ‘Cooperation and cheating in *Pseudomonas aeruginosa*: the roles of the *las*, *rhl* and *pqs* quorum-sensing systems’, The ISME Journal. Nature Publishing Group, 5(8), pp. 1332–1343. doi: 10.1038/ismej.2011.13.

Williams, P. and Cámara, M. (2009) ‘Quorum sensing and environmental adaptation in *Pseudomonas aeruginosa*: a tale of regulatory networks and multifunctional signal molecules’, Current Opinion in Microbiology, 12(2), pp. 182–191. doi: 10.1016/j.mib.2009.01.005.

Winstanley, C., O’Brien, S. and Brockhurst, M. A. (2016) ‘*Pseudomonas aeruginosa* Evolutionary Adaptation and Diversification in Cystic Fibrosis Chronic Lung Infections.’, Trends in microbiology. Elsevier, 24(5), pp. 327–337. doi: 10.1016/j.tim.2016.01.008.

## Supplementary References

Bao, Y. et al. (1991) ‘An improved Tn7-based system for the single-copy insertion of cloned genes into chromosomes of gram-negative bacteria’, Gene. Gene, 109(1), pp. 167–168. doi: 10.1016/0378-1119(91)90604-A.

Jayakumar, P. et al. (2021) ‘Pseudomonas aeruginosa reaches collective decisions via transient segregation of quorum sensing activities across cells’, bioRxiv, p. 2021.03.22.436499. doi: 10.1101/2021.03.22.436499.

De Lorenzo, V. et al. (1990) ‘Mini-Tn5 transposoon derivatives for insertion mutagenesis, promoter probing, and chromosomal insertion of cloned DNA in gram-negative eubacteria’, Journal of Bacteriology. American Society for Microbiology Journals, 172(11), pp. 6568–6572. doi: 10.1128/jb.172.11.6568-6572.1990.

